# Frontal P3 Potential as a Supramodal Marker of Imminent Attentional Lapses

**DOI:** 10.64898/2026.05.20.726475

**Authors:** J.L. Kenemans, E. Canny, J. van der Haest, D. Koevoet

## Abstract

Focusing on an organism’s task at hand is instrumental for intelligent and goal-driven behavior. However, humans and other animals often fail to pay sustained attention across long time intervals. Failing to stay on-task may cause one to miss crucial task-relevant signals, leading to impaired performance, which can have serious consequences. Therefore, it is important to understand the neural basis of attentional lapses. One promising neural marker of attentional lapses is the frontal P3 (fP3) EEG component, which has been suggested to reflect the susceptibility to incoming sensory input. Following this, we hypothesized that the fP3 1) predicts imminent lapses of attention, and 2) that it should predict upcoming lapses of attention across modalities. In two experiments, we found that the fP3 reliably tracked lapses of attention of sustained attention already seconds preceding the crucial visual signal. We further extended this to the auditory domain: Already 1.5s ahead of the incoming auditory target, the fP3 revealed whether that target was detected or not. Detailed topographic analyses did, however, reveal a slight dissociation between modalities in underlying intracranial source configurations. In sum, this work revealed a supramodal neural signature of susceptibility, which tracks lapses of sustained attention seconds ahead of the critical incoming sensory input.

## Introduction

Focusing on one’s task at hand is a cornerstone of intelligent and adaptive behavior (Nobre & Coull, 2010). Yet, humans and other animals often fail to pay sustained attention throughout longer time intervals. For example, when driving for many hours without taking a break, one may miss crucial incoming sensory information, potentially leading to serious consequences. In addition to such applied implications, fundamental scientific questions of how sensory information is processed based on the brain’s state serve as strong motivations to uncover the neural basis of lapses of sustained attention.

In a series of older and more recent studies it has been demonstrated that in certain cognitively demanding situations the human electro-cortical response to relatively unexpected and salient external signals is reduced. This reduction has been observed especially for simulated car-driving conditions (Scheer, Bülthoff, & Chuang, 2016; Van der Heiden et al., 2018; Wester, Bocker, Volkerts, Verster, & Kenemans, 2008), autonomous driving (Van der Heiden et al., 2018), but also cognitive load as induced by language production (Janssen, Schutte, & Kenemans, 2024; Van der Heiden, Janssen, Donker, & Kenemans, 2020; Van der Heiden, Kenemans, Donker, & Janssen, 2022). Note that in all these studies the electro-cortical response was elicited by task-irrelevant stimuli.

The reduced electro-cortical response in these studies concerned an EEG event-related potential (ERP) component elicited by task-irrelevant auditory, unique, infrequently presented, salient, “novels” (e.g., honking car, barking dog). This so-called frontal P3 (fP3) has a peak latency of about 300ms post-novel and has been postulated to reflect a transient generalized state of neural and motoric inhibition, implemented in or originating from dorsalmedial frontal cortex (Kenemans, 2015; Polich, 2007; Wessel & Aron, 2013, 2017). Independent of this association with neural and motoric inhibition, fP3 has also been suggested as a measure of susceptibility to salient, potentially relevant external signals outside the primary focus of attention. Following Van der Heiden et al. (2022, p.1196), “susceptibility refers to the extent to which an observer is in a mode that allows for detection of external signals to such a degree that an adequate behavioral response can be based on (…) detection.”

However, while the above studies demonstrated that an fP3 is elicited by irrelevant auditory stimuli in a visual-task context, they do not provide direct evidence for an association between fP3 and susceptibility. Fortunately, another strain of work demonstrated that visually-evoked fP3s are predictive of upcoming successful or failed detection of visual signals. These studies employed the continuous temporal expectancy task (CTET), wherein a checkerboard-like stimulus rotates continuously every 800ms. Participants are tasked to detect whenever the stimulus remains in a specific rotation for a slightly longer duration. O’Connell et al. (2009) found that an fP3 elicited by rotating checkerboard stimuli was reduced when a subsequent visual target was missed, versus when it was correctly detected (Chidharom, Krieg, Marques-Carneiro, Pham, & Bonnefond, 2021; Dockree et al., 2017; also see Martel, Dähne, & Blankertz, 2014).

In the present work we aimed to (1) provide another replication of this prospective association, (2) detail its temporal profile with substantially more statistical power than previous work, and (3) extend this to the auditory domain, to provide support for the notion of fP3 reflecting auditory susceptibility.

To this end, we conducted two EEG experiments wherein we directly tested the prospective association between the fP3 and imminent lapses of sustained attention. Previewing our results, we found upcoming hits were associated with larger fP3 amplitudes than misses did already seconds prior to the crucial visual input, replicating previous work. In our second experiment, we found that the fP3 served as a supramodal marker of imminent lapses of sustained attention: in both the visual and auditory versions of the task we found larger fP3 amplitudes preceding upcoming hits compared with misses. Lastly, we conducted a detailed topographical comparison between the visual and auditory fP3 to determine the extent of overlap between their intracranial equivalent-dipole configurations.

## Method Experiment 1

### Participants

Data were collected for a total of 27 participants, which exceeds prior work (O’ Connell et al., 2009). All participants reported no current or previous psychiatric diagnoses or neurological disorders, to not be smokers (no cigarettes in the last 6 months), normal or corrected-to-normal vision and no history of epilepsy (in relation to flickering of stimuli during one of the tasks). Additionally, participants were asked to not consume coffee, alcohol, stimulants or other drugs starting from 10PM the evening before participation. Two participant data sets were discarded because less than 10 trials for either hits and/or misses were retained after artifact rejections and blink-on-target detection, leaving a final sample of 25 (*M*_age_= 22.6, range = [20-32], 22 females, 3 males). The experiment was approved by the ethical review board of the Utrecht University Faculty of Social Sciences.

### CTET

Throughout the continuous temporary expectancy task (CTET; Figure 1), participants were presented with a continuous stream of ‘frames’. A single frame consisted of a centered 10×10 grid of 8mm^2^ squares split into black and white halves on a gray background (see O’Connell et al., 2009); the full frame extended across 8×8 dva. Frame orientation randomly turned 90^°^ clockwise or counterclockwise every frame presentation (resulting in four different grids). To limit eye movements, a white fixation cross was presented throughout the task. In addition, stimuli flickered at a constant rate of 25Hz.

**Figure 1.**
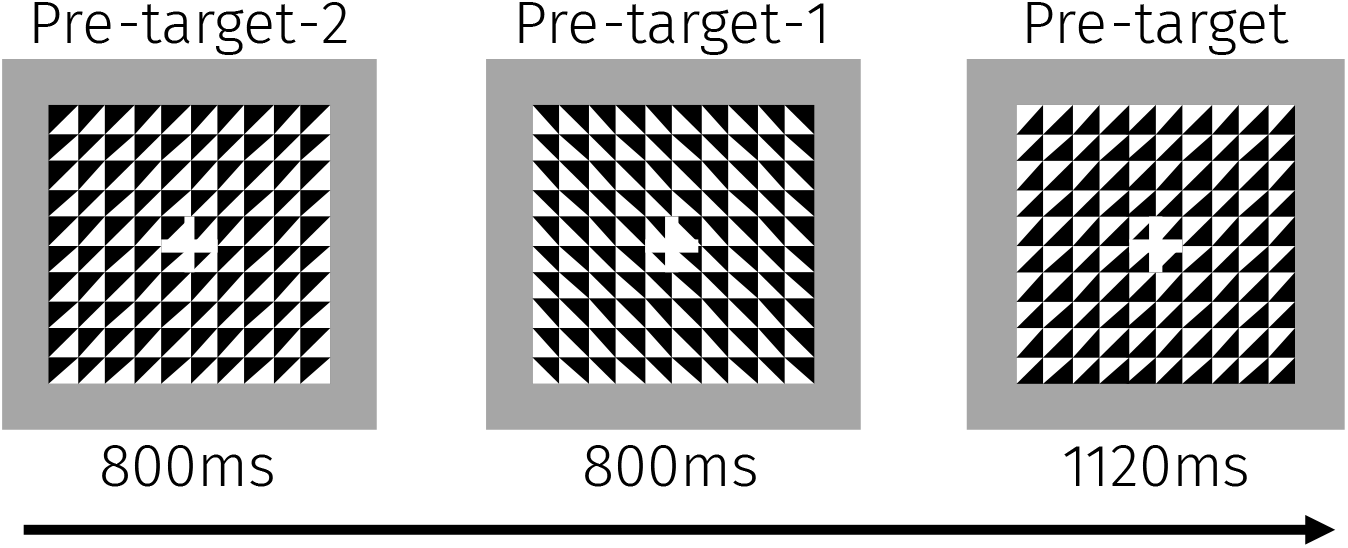
Task design of the visual continuous temporary expectancy task (CTET). Participants detected frames that were presented 1120ms among non-target frames that were presented only 800ms.

Participants were seated 57cm from the monitor and were instructed to identify target frames, which had an increased duration (1120ms) compared to non-target frames (800ms). This implies that targets could be distinguished from non-targets after 800ms of target presentation – this timepoint will be referred to as time of discrimination (ToD). To identify a target, participants were instructed to press the spacebar key as fast as possible on a keyboard with their dominant index finger. Stimuli were presented in pseudo-random order so that 7-15 non-target frames (average of 11 frames; corresponds to 5.6s-12s, average of 8.8s) were presented between target frames. Each block consisted of 255 frames and lasted for approximately 3 minutes and 25s. The number of target frames across blocks varied between 199-213, with a mean of 206 target frames across the full experimental session. Subjects completed 10 blocks with small rest breaks (∼1 min) in between blocks.

Participants completed two practice rounds consisting of two blocks. Target frame presentation was increased to 1280ms during practice so that participants could establish a solid understanding of the task. In the first practice block, subjects had to identify three target frames randomly interspersed among 25 non-target frames. In this block the 25Hz flicker was omitted. The second practice block was identical to the first, but now the 25Hz flicker was included. Both blocks were completed twice.

### EEG recording

EEG was recorded with the ActiveTwo system (Biosemi, Amsterdam, The Netherlands) using 64 Ag/AgCl electrodes placed according to the international 10/10 system. Vertical electrooculogram (EOG) was recorded from electrodes positioned above and below the left eye. Horizontal EOG was recorded from electrodes placed at the outer canthi of both eyes. Recorded signals were online referenced to the Common Mode Sense/Driven Right Leg electrodes, and the sample rate was 2048Hz with a low-pass filter of 417Hz. Electrodes were also placed on the left and right mastoids for later offline re-referencing.

### Data analysis

Spacebar presses were counted as hits when they occurred in a time window from 150 to 1600ms after ToD (or 950 to 2720ms after onset of a pre-target (PT) frame, a frame that after 800ms turned into a target; cf. Dockree et al., 2017). False-alarm rates were calculated as the number of spacebar presses outside a target window, divided by the number of non-target frames.

Using Brain Vision analyzer (BVA) 2.1, EEG data were re-referenced to the average mastoid signals; down-sampled to 256 Hz (after application of an anti-aliasing filter); low-pass filtered at 40 Hz; high-pass at 0.5 Hz; and subjected to a notch filter at 50 Hz. Subsequently EEG data were segmented into epochs from -4000ms to 1700ms relative to PT onset.

We labeled epochs as containing an artifact if 1) a change in voltage exceeded 50 µV/ms; or 2) changes in voltage were lower than a change of 0.5 µV/100ms; or 3) absolute amplitude exceeded 200 µV – all within a time window of -4000 to 500ms relative to PT onset. The influence of ocular artifacts was attenuated using the BVA implementation of the Gratton-Coles algorithm. We excluded all epochs labeled as artifact trials from all analyses. Additionally, we excluded epochs wherein participants blinked during the presentation of the target frame. We detected such epochs by examining when changes in VEOG amplitudes exceeded 200 µV between -500 and 1700ms relative to PT onset. As mentioned, participants with fewer trials than 10 for either hits or misses were excluded.

Finally, epochs lasting from -4000 to 500 relative to PT onset were averaged separately for hits and misses to obtain multi-frame average ERPs aligned to PT and 4 preceding non-target frames, termed PT-1 (immediately preceding PT), PT-2 (2 back), PT-3 (3 back), and PT-4 (4 back). These average ERPs were additionally filtered using a 25-Hz band rejection (to reduce the SSVEP), and an 8-Hz low pass setting (to reduce residual noise).

### fP3 analysis

Frontal P3s were identified in the post-PT epoch, after baseline correction by subtracting the average values across 80ms preceding PT onset. Figure 2 shows the resulting grand-average PT scalp distributions across 25 participants in a time window (280-400ms) where the peak fP3 could be expected (O’Connell et al., 2009). Following O’Connell et al., we first set out to define a pre-component baseline (usually centered around the negative peak preceding the positive fP3). Secondly, given this baseline, fP3 itself was identified using cluster-based logic. Thirdly, the pre-component baseline as well as the latency windows identified with cluster-based logic were used to extract fP3 values for PT, as well as for PT-x, 1<=x<=4. These steps are detailed right below.

**Figure 2.**
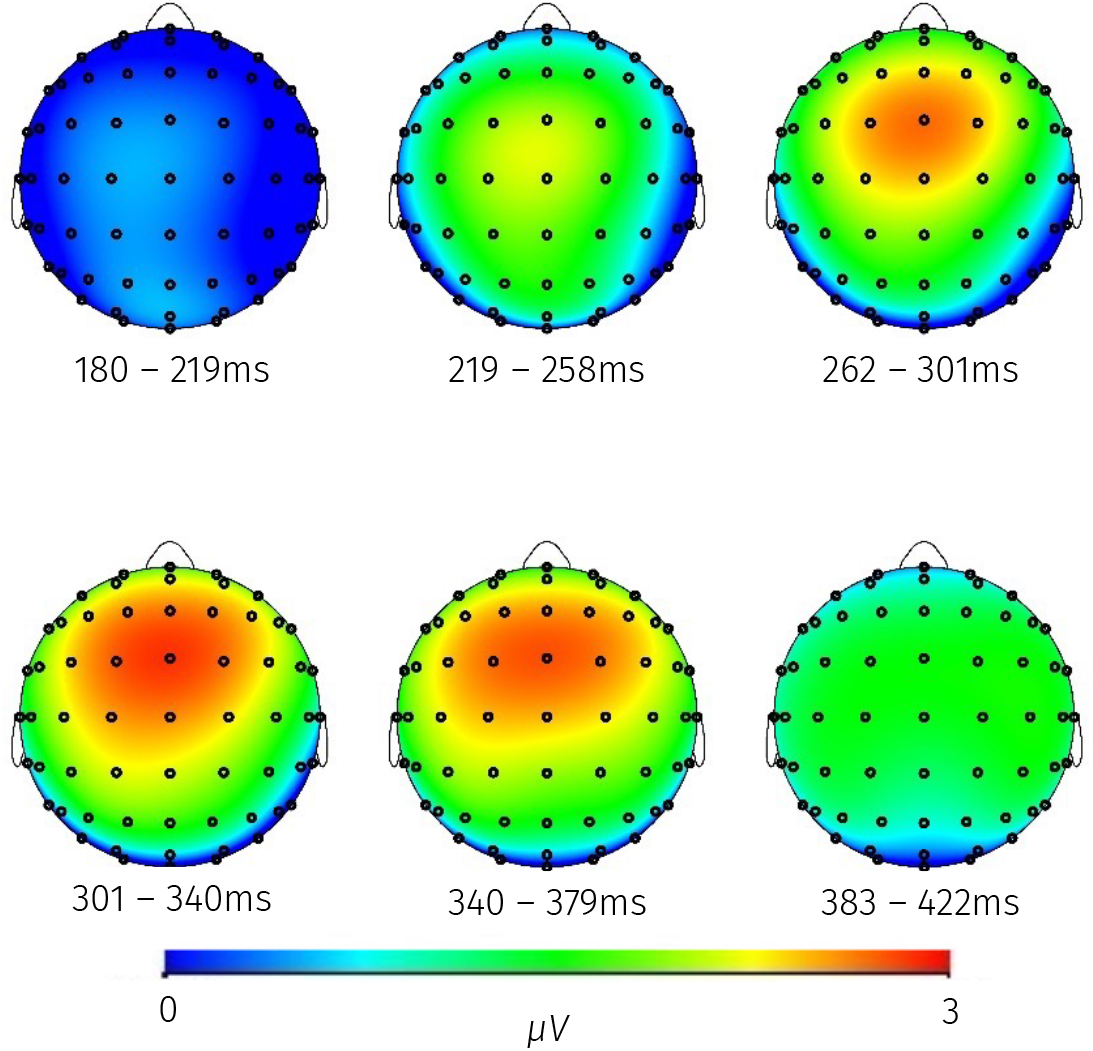
Grand-average scalp distributions in the expected post-PT time window for fP3. Average across hits and misses. Baseline -80 to 0ms preceding the pre-target interval.

As can be seen in Figure 2, the peak value centers around the FCz location between 300 and 340ms. Following O’Connell et al. a pre-component baseline was then selected. From the waveform at FCz the grand-average peak of the preceding negative peak was detected between half-maxima at 114 and 176ms; consequently, the pre-component baseline interval was chosen as 114-176ms and the average amplitudes in that interval were used for baseline correction.

We subsequently used a cluster-based approach to detect time points where the fP3 was larger for hits than for misses (for the PT exclusively), as in Chen, van de Vijver, Canny, Kenemans, and Baas (2024). Because the focus was on only one sensor (FCz) and involved 66 baseline samples (-80 to 176ms) and 66 active samples (180 to 434ms), critical alpha values were set equal for the baseline and the active period; therefore active clusters of significance (minimal length 2 samples) contained only *p* values for the Hit-Miss difference that were smaller than the smallest baseline *p* value. This yielded two clusters of significance: 180-231ms (all *p*s < .04) and 278-332ms (all *p*s < .04).

Finally, the 180-231 and 278-332 windows were used to create average amplitudes for each subject and Hit/ Miss condition at FCz, for PT, and for PT-1 to PT-4. In all cases the average value in the 114-176ms baseline interval was subtracted from these average amplitudes.

Throughout the paper, we used a multivariate testing procedure (SPSS v31). This approach is similar to repeated-measures ANOVA but has the advantage of not assuming sphericity and ensures that the total number of degrees of freedom does not exceed the total number of independent observations (i.e., sample size). Note that when using repeated-measures ANOVAs we obtained highly similar results (i.e., same significances and highly similar effect sizes; data not reported).

## Results and Discussion Experiment 1

### Performance

Average hit rate was 59.3% and average false-alarm rate was 1.3%. Mean response time for hits was 556.9ms (*SEM* = 16.78ms).

### fP3

Figure 3 shows the waveforms for PT-4 to PT-1 and PT. Prospective associations were further analyzed with a 2 Latency (early vs. late cluster) x 5 Pre-target interval (PT, PT-1, 2, 3, 4) x 2 Detection (hit vs. miss) multivariate repeated-measures ANOVA (Supplementary Table 1). We observed larger fP3 amplitudes prior to a hit compared with a miss (*F*(1,24) = 10.75, *p* = .003,*η*_*p*_^*2*^ = .31). This indicates that smaller fP3 amplitudes predicted upcoming lapses of attention. Regarding the time course of this effect, we found no significant interactions of Detection with Pre-target interval or with Latency (*F*s < 1, *p*s > .42, *η*_*p*_^*2*^s < .17).

**Figure 3.**
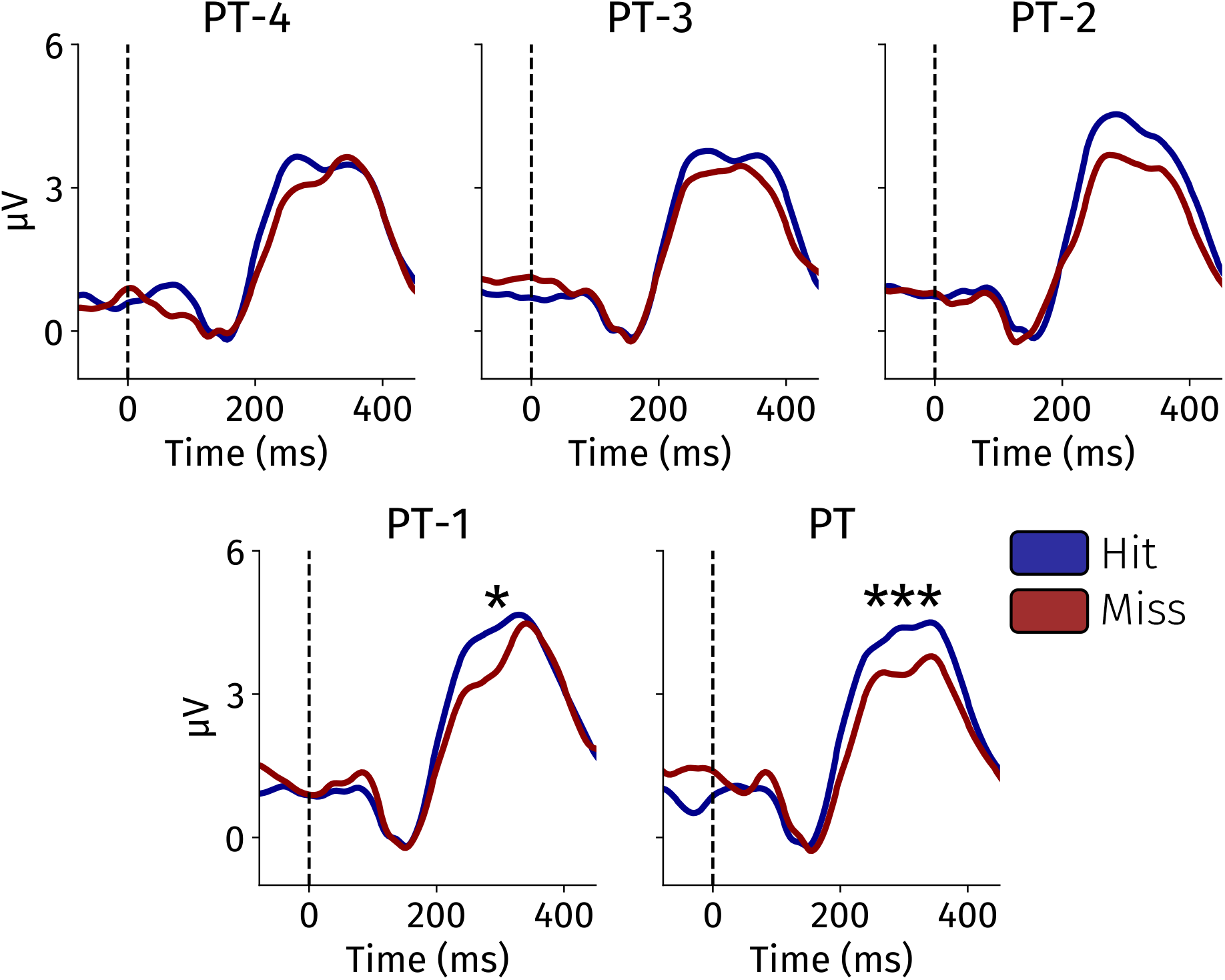
Waveforms for FCz, hits (blue) and misses (red). From upper left to lower right: PT-4 to PT. Note zero values between 114 and 176ms due to pre-component baseline correction.*** Hit/Miss difference significant at p < .005, for PT pooled with PT-x (x = 1,2,3,4). * Hit/Miss difference significant at p < .05, for PT-x pooled with PT-y (y>x).

Despite the non-significant effect interaction effect between Detection and Pre-target interval, we wanted to examine *when* the fP3 started to reveal imminent lapses of attention. This is important to consider from a brain-behavior-relationship perspective but also because the timing of this effect is vital to potential applications. We examined this issue through a stepwise removal approach. That is, we iteratively removed the PT closest in time to the imminent hit or miss and reconducted the repeated-measures ANOVA as above. We stopped this procedure whenever the difference in fP3 amplitude no longer significantly differed between hits and misses. This stepwise removal approach allowed us to further characterize the timing of the Detection effect while maximizing statistical power. Turning to the results, we still observed a significant Detection effect when removing PT from the analysis (*F*(1,24) = 5.40, *p* = .023, *η*_*p*_^*2*^ = .18), but not when removing PT and PT-1 from the analysis (*F*(1,24) = 3.81, *p* = .063, *η*_*p*_^*2*^ = .14). This indicates that fP3 amplitudes were prospectively associated with upcoming lapses of attention approximately 1.5s before the crucial input signal.

The above results indicate that fP3 amplitudes reliably track upcoming lapses of attention. We found this effect to be reliable already approximately 1.5s before target detection was possible. In line with previous work (O’Connell et al., 2009), this indicates that the fP3 may reflect the susceptibility to incoming sensory input, which reliably predicts upcoming lapses of attention. We pushed this further in Experiment 2. First, we nearly doubled our sample size to increase statistical power so that we could examine the robustness of our findings. This also allowed us to more closely examine *when* the fP3 started to predict an upcoming lapse of attention. Second, as the fP3 has been suggested to reflect susceptibility to both visual and auditory inputs, we here tested whether the fP3 could also predict upcoming auditory lapses of attention.

## Method Experiment 2

### Participants

In Experiment 1, the sample size was based on earlier literature and may have resulted in insufficient power to detect prospective associations earlier than 1500ms preceding ToD. Furthermore, we had no detailed estimates of effect size regarding the auditory CTET. Therefore, it was decided to aim for a doubling of the sample size relative to experiment 1 (*n=* 50). Eventually, data were collected for a total of 48 participants. All participants reported no current or previous psychiatric diagnoses or neurological disorders, to not be smokers (no cigarettes in the last 6 months), normal or corrected-to-normal vision and no history of epilepsy (in relation to flickering of stimuli during one of the tasks). Additionally, participants were asked to not consume coffee, alcohol, stimulants or other drugs starting from 10PM the evening before participation. Although we slightly undershot our target sample of 50 participants due to practical limitations, we still included nearly double the number of participants as in Experiment 1.

For the visual CTET, one participant-data set was lost due to disk-saving failure, and for another data collection was halted due to insufficient performance during practice. In total, 46 participants were included in these analyses (*M*_age_ = 23.2, range = [18-36], 27 females, 19 males). For the auditory CTET four participant-data sets were discarded because these participants were unable to sufficiently discriminate the pitches for adequate performance; one data set was incomplete due to disk-saving failure; and three further participants produced less than 10 misses. We included 40 participants in our analyses (*M*_age_ = 23.4, range= [18-36], 24 females, 16 males). For a total of 39 participants, both visual and auditory data were available (*M*_age_ = 23.4, range = [18-36], 23 females, 16 males). No participant-datasets were discarded because of less than 10 trials for either hits and/or misses after artifact rejection. The experimental procedures were approved by the ethical review board of the Utrecht University Faculty of social Sciences.

### CTET

Participants completed the visual and auditory versions of the CTET. The order of tasks was counterbalanced across participants.

The visual CTET was identical to the procedure from Experiment 1 (Figure 1), except for the 25 Hz flicker, which was maintained for only 10 participants; for 27 participants the flicker was at 12 Hz, while for the remaining 9 there was no flicker. Although there is no reason to expect an interaction between flicker-characteristic effects and other effects on fP3, we nevertheless conducted a post-hoc statistical analysis to test this potential interaction.

We based our version of the auditory CTET on a previous behavioral study (Berry, Li, Lin, & Lustig, 2014). Throughout the auditory CTET (Figure 4A), participants were exposed to a continuous stream of sine wave tones played through earbuds (ER3C Insert Earphones, Etymotic Research) at an initial sound pressure of 70 dB, while viewing a white fixation cross on a screen with a grey background to limit eye movements. Because of complaints about the intensity, for the final 7 participants the loudness was reduced to 50 dB. Four different pure-tone pitches (500, 625, 750, 875 Hz) alternated randomly with 800ms duration each; there was no gap between successive tones. Target tones lasted 970ms instead of 800ms, to produce performance comparable to that in the visual CTET (based on pilot data). Further stimulus-configuration parameters were identical to those in the visual CTET.

**Figure 4.**
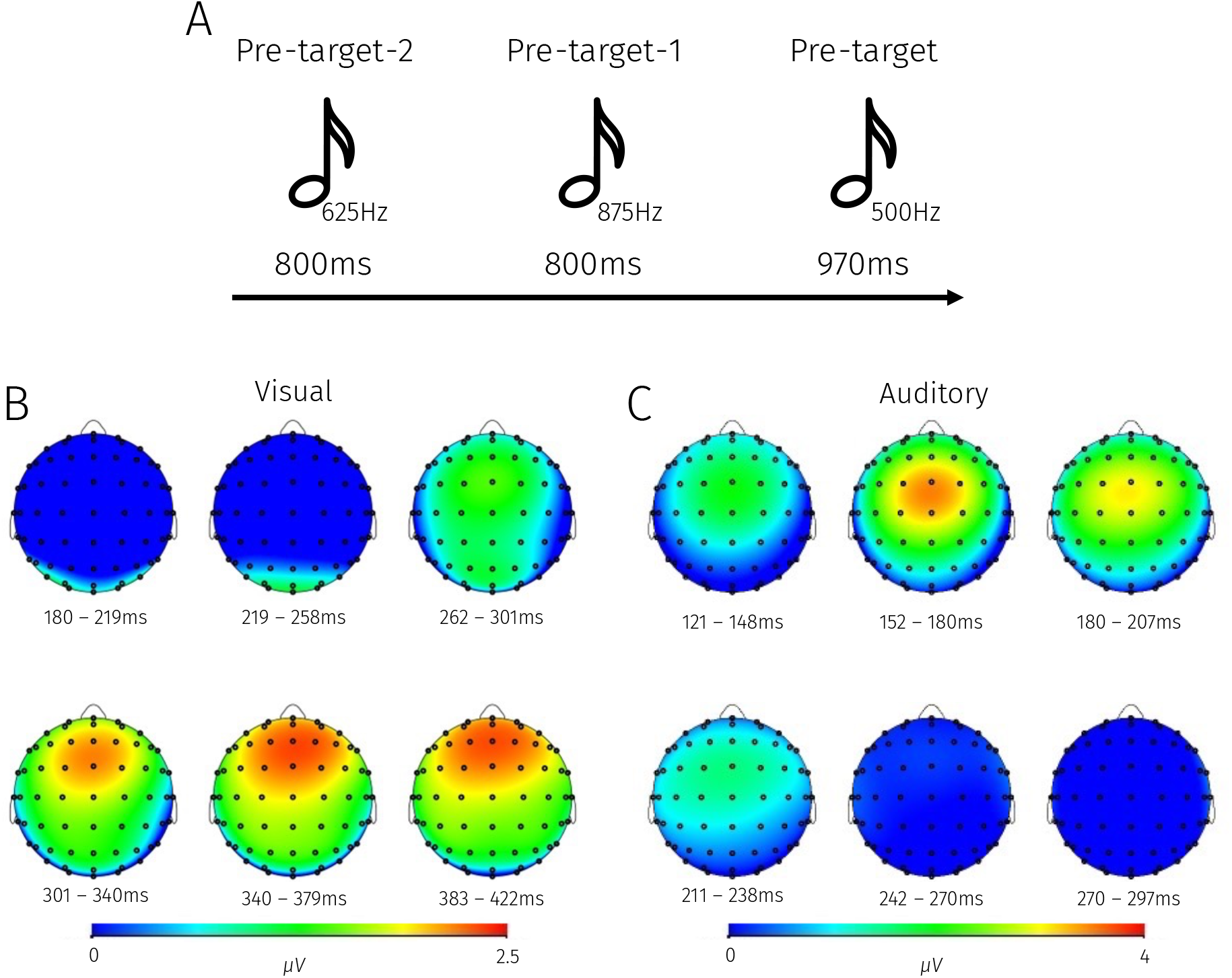
Auditory CTET task design and grand-average scalp distributions for Experiment 2. **A** Participants detected target tones that were presented for a longer duration (970ms) compared with shorter duration non-target tones (800ms). **B** Scalp topographies computed during the PT interval for visual (left) and **C** auditory (right) versions of the task. All topographies reflect average amplitudes across hits and misses. Baseline -80 to 0ms preceding the pre-target.

### Data recording and analysis

EEG recording and performance analysis followed the same procedures as in Experiment 1. This also held for EEG-data processing. The only difference was that we did not remove trials with blinks in the (pre-)target time window for auditory CTET data.

### fP3 analysis

For the visual CTET, Figure 4B shows the grand-average pre-target (PT) scalp distributions across 46 participants in a time window (280-400ms) where the peak fP3 could be expected. Based on the results of Experiment 1, fP3 amplitude was quantified as the average amplitude at FCz across latency windows 180-231ms and 278-332ms, separately for each participant, Hit/ Miss condition, and PT frame. For baseline correction, again the average value in the latency window 114-176ms was used.

The auditory fP3 effect was expected to be earlier or at the same latency as visual fP3 (e.g., Kenemans, 2025; Kenemans et al., 2023). As can be seen in Figure 4C, the peak value centers around the FCz location between 150 and 200ms. From the waveform at FCz the grandaverage peak of the preceding negative peak was detected between half-maxima at 86 and 113ms; to obtain a baseline interval of a width comparable to previous studies (e.g., Kenemans et al., 2023) the time window 63-113ms was chosen, and the average amplitudes in that interval were used for baseline correction. Subsequently cluster-based logic was again applied to detect time points where PT fP3 was larger for hits than for misses (Chen et al., 2024). The cluster-based analysis involved 50 baseline samples (-80 to 113ms) and 50 active samples (117 to 300ms). Critical alpha values were set equal for the baseline and the active period; therefore, active clusters of significance (minimal length 2 samples) contained only *p*values for the Hit-Miss difference that were smaller than the smallest baseline *p*-value. This was found for a time window of 129 to 207ms; note however that nowhere in this latency range *p*-values were lower than 0.11. The values in this latency range were used to create average amplitudes for each participant and Hit/ Miss condition at FCz, for PT and for PT-1 to PT-4.

## Results Experiment 2, Visual CTET

Average hit rate was 49.4 % and average false-alarm rate was 2.1%. Mean response time for hits was 698.6ms (*SEM* = 19.0ms).

Figure 5 shows the waveforms for PT-4 to PT-1 and PT. Prospective associations were examined with a multivariate repeated-measures ANOVA using Latency (180-231 vs. 278-332) x Pre-target interval (PT, PT-1, 2, 3, 4) x Detection (Hit vs. Miss) as predictors (Supplementary Table 2). fP3 amplitudes were significantly larger prior to upcoming hits compared with misses (*F*(1, 45) = 26.45, *p* < .001, *η*_*p*_^*2*^ = .37). This effect differed between latencies (*F*(1, 45) = 14.47, *p* < .001, *η*_*p*_^*2*^ = .24), Specifically, the difference between hits and misses was more pronounced for the longer (*F*(1,45) = 32.38, *p* < .001, *η*_*p*_^*2*^ = .42) than for the shorter latency (*F*(1,45) = 9.70, *p* = .003, *η*_*p*_^*2*^ = .18), but remained significant in both time windows.

**Figure 5.**
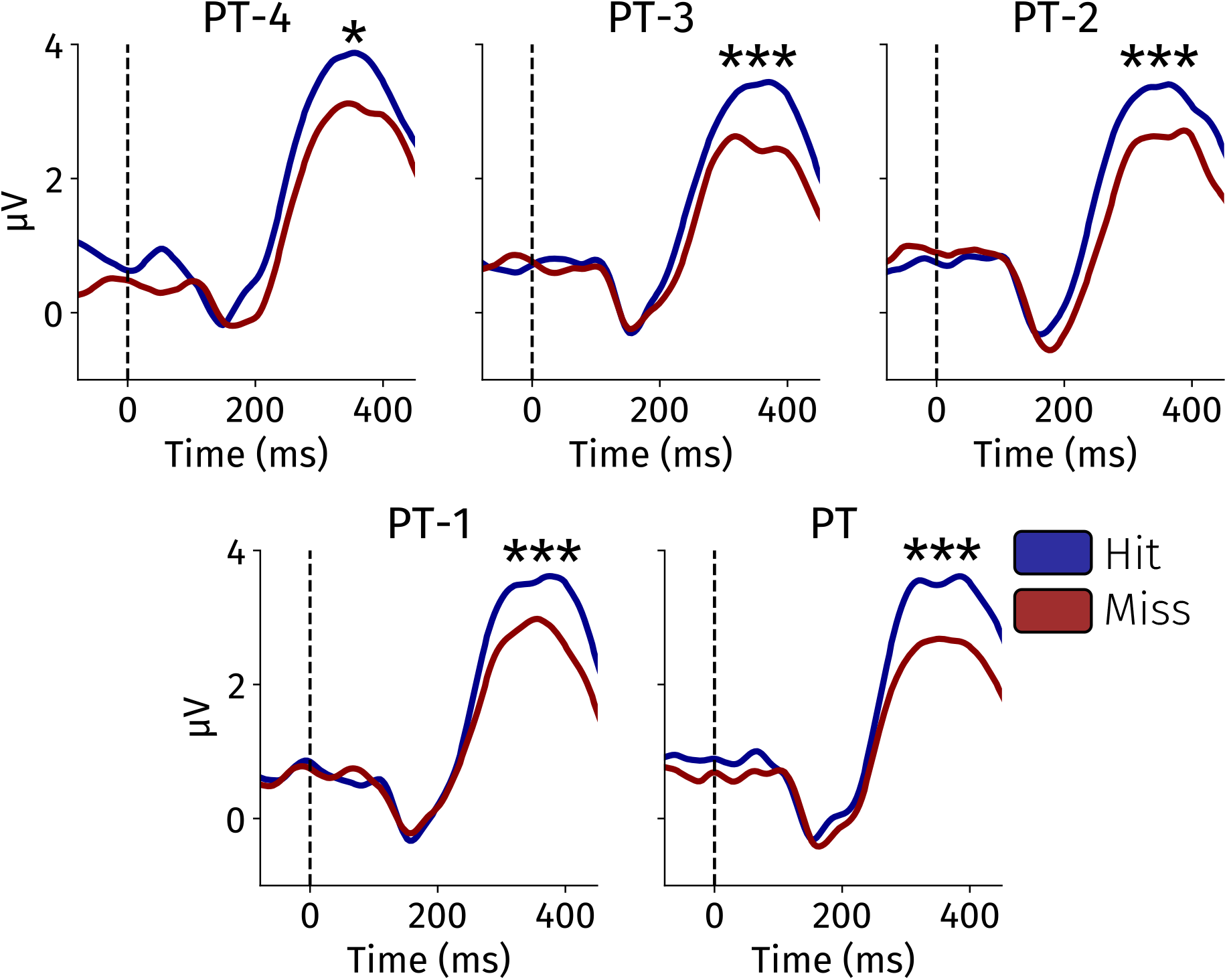
Waveforms for FCz in the visual CTET, plotted separately for imminent hits (blue) and misses (red). From upper left to lower right: PT-4 to PT. Note zero values between 114 and 176ms due to pre-component baseline correction. *** Hit/Miss difference significant at p < .001 for PT pooled with PT-x and PT-x pooled with PT-y (x = 1,2,3,4; y>x). * Hit/Miss difference significant at p < .05, for PT-x pooled with PT-y (y>x).

As the Detection effect was strongest for the longer latency, we examined *when* the fP3 differed between imminent hit and miss trials for this latency. To this end, we removed one of the frames in a stepwise fashion and reconducted the repeated-measures ANOVA (as in Experiment 1). Iteratively removing PT, PT-1, PT-2, and PT-4 did not remove the significant detection main effect at any step (all *F*s(1,45) > 5.75, *p*s < .03, *η*_*p*_^*2*^s > .11; although note that the effect was numerically weaker when analyzing PT-4 in isolation; Figure 4). This demonstrated that fP3 amplitudes were prospectively associated with imminent lapses of attention already ∼4s before the incoming crucial input.

Since we collapsed data across different flicker conditions in Experiment 2, we repeated the main analysis above while also including the flicker condition (i.e., no flicker, 12 Hz, and 25 Hz flicker) in our model as a between-subjects factor. This allowed us to examine whether different flicker conditions drove our effects of interest. Detection and Latency x Detection effects were again highly significant (both *p* < .006), without any effect of Pre-target interval. Importantly, neither of these effects were modulated by flicker condition (all *p*s > .3, all η_*p*_^*2*^s < .068). Overall, fP3 was slightly smaller for 12-Hz flicker as compared to no or 25-Hz flicker (*F*(2,68) = 4.2, *p* < .05, *η*_*p*_^*2*^=.11). Mean fP3s for none, 12, and 25 Hz were 0.8, 0.3, and 1.2 (short latency), and 3.8, 2.7, and 3.4 µV (long latency), respectively.

In this larger sample we also examined the possibility of a spurious correlation between fP3 and target detection. It is conceivable that with longer non-target sequences expectancy for the target builds up, resulting in larger fP3s to non-targets and an increased detection probability. This would also result in misses being preceded by smaller overall fP3s compared to hits, which should manifest in an increasing fP3 when proceeding from PT-4 to PT, irrespective of whether a hit or a miss subsequently ensues. However, we found no main effect of the pre-target interval (*F*(4,42) = 1.53, *p* = .14, *η*_*p*_^*2*^ = .13), nor an interaction between detection and pre-target interval (*F*(4,42) = 0.42, *p* = .79, *η*_*p*_^*2*^ = .04), which rules out this possible alternative explanation for our results (also see Figure 5).

As in Experiment 1, we found that fP3 amplitudes predicted upcoming lapses of sustained attention. Equipped with higher statistical power, we were able to uncover that fP3 responses to pre-target frames occurring up to 4 seconds preceding possible target detections tracked attentional lapses. Our data demonstrate that the fP3 is a sensitive and early marker of upcoming lapses to incoming visual inputs.

## Results Experiment 2, Auditory CTET

Turning to the auditory CTET data, average hit (49.1%) and false-alarm rates (3.1%) were comparable to that for the visual CTET. Response times for hits were on average 727.15ms (*SEM* = 19.1ms). In the 39 participants who completed both the visual and auditory tasks, we found no significant difference in hit (*t*(38) = 1.18, *p* = .24, *d* = .19) and false-alarm rates (*t*(38) = 1.15, *p* = .26, *d* = .18) between modalities.

Figure 6 shows the waveforms for PT-4 to PT-1 and PT. Again, fP3 amplitudes were submitted to a multivariate repeated-measures ANOVA (Supplementary Table 3), 5 (PT, PT-1, 2, 3, 4) x 2 (Hit vs. Miss) design. Extending our findings from the visual to the auditory domain, we found larger fP3 amplitudes for upcoming hits compared with misses (*F*(1,39) = 6.17, *p* = .017, *η*_*p*_^*2*^= .14). As in our previous analyses, we found no significant interaction effect between pretarget frame number and detection condition (*F*(4,36) = 1.03, *p* = .406, *η*_*p*_^*2*^ = .10).

**Figure 6.**
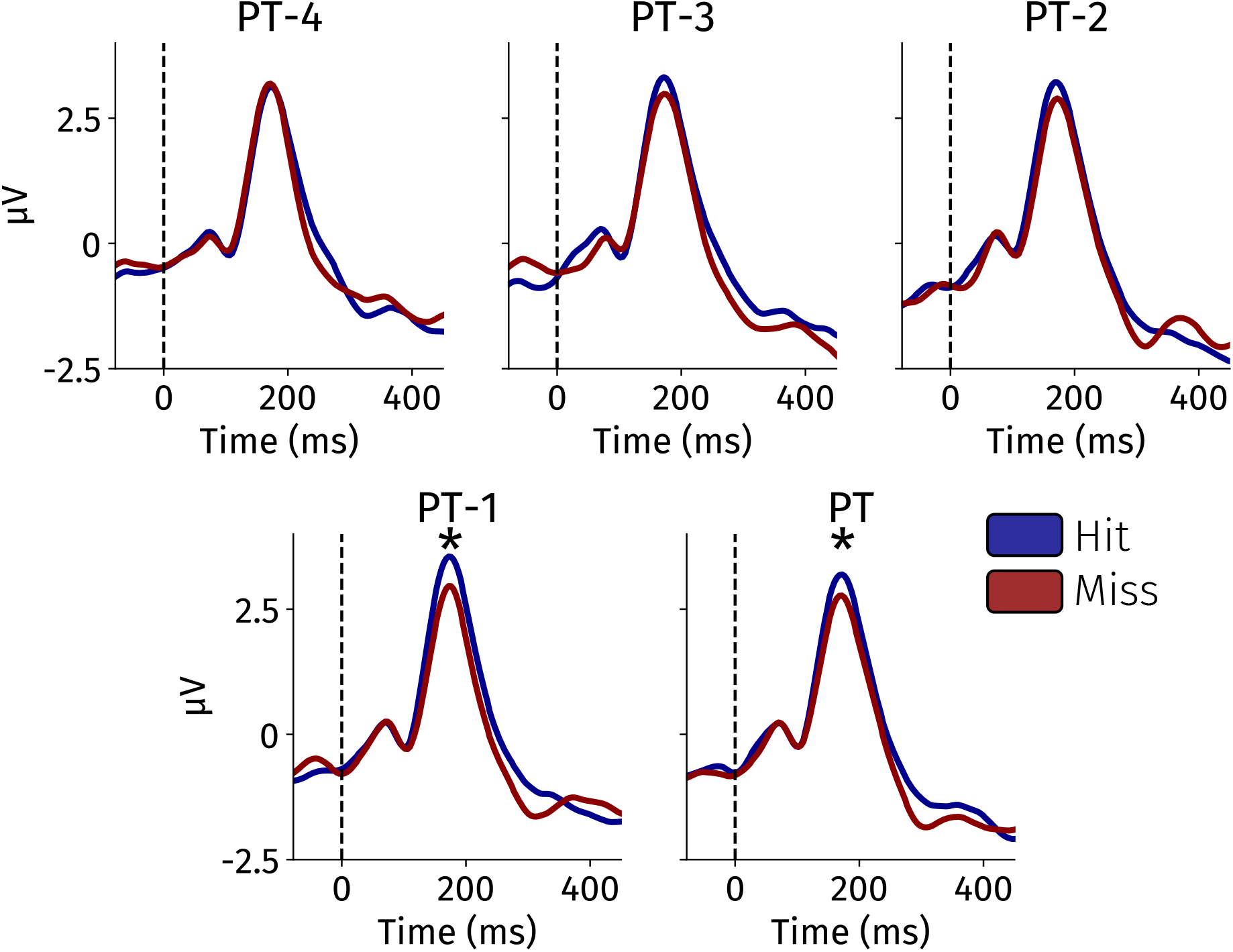
Waveforms for FCz in the auditory CTET, plotted separately for imminent hits (blue) and misses (red). From left to right: PT-4 to PT. Note zero values between 63 and 113ms due to precomponent baseline correction. * Hit/Miss difference significant at p ≤ .05, for PT-x pooled with PT-y (y>x).

**Figure 6.**
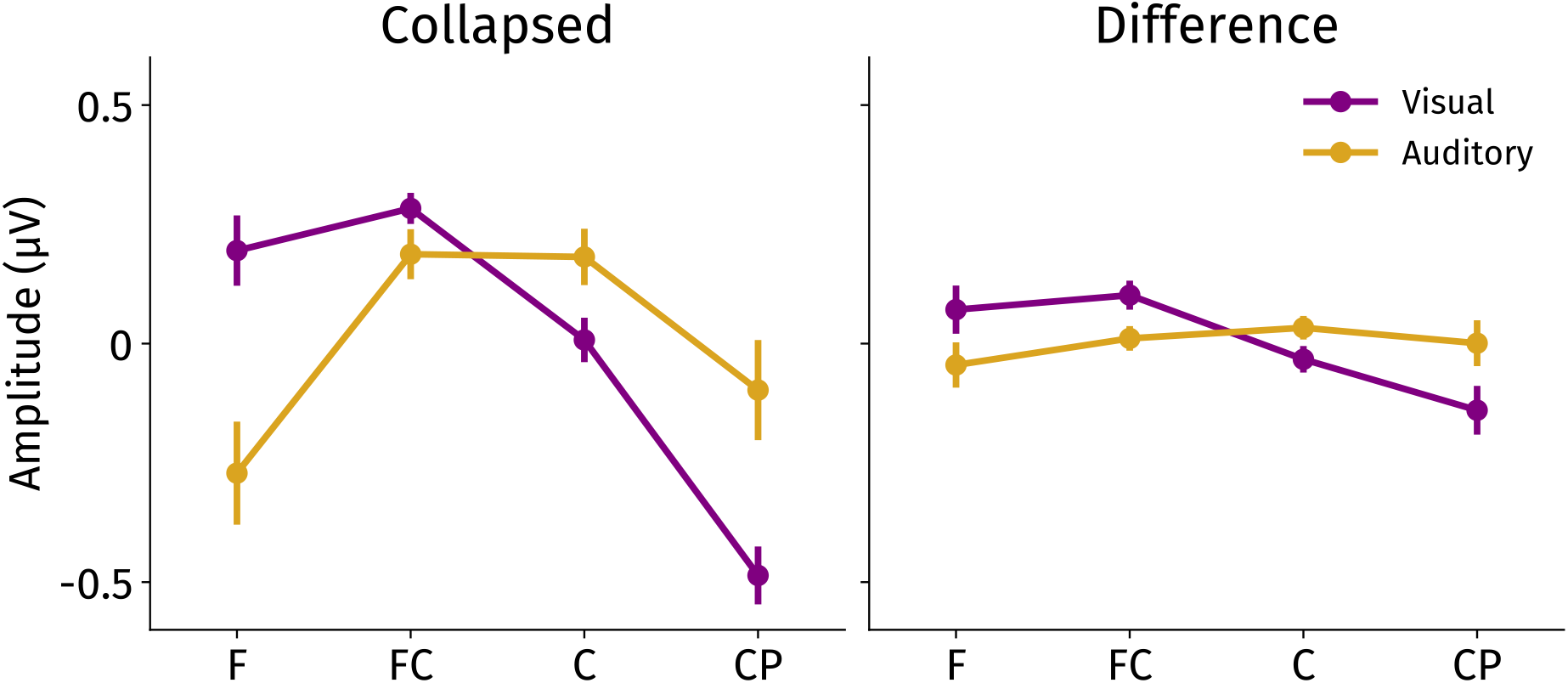
Left: Average-referenced fP3 amplitudes as a function of Modality and Anteriority collapsed across hits and misses. Right: Same for the difference between hits and misses. Error bars reflect standard errors of the mean. F = Frontal; FC = Frontal-Central; C = Central; CP = Central-Parietal.

As before, we performed a stepwise removal analysis to examine *when* the fP3 started to become associated with imminent lapses of attention. When removing PT, the detection effect was on the threshold of significance (*F*(1,39) = 4.09, *p* = .050, *η*_*p*_^*2*^= .10). Upon removal of PT-1, the detection effect was no longer significant (*F*(1,39) = 1.10, *p* = .301, *η*_*p*_^*2*^ = .03). This demonstrated that the fP3 was associated with upcoming lapses of attention ∼1-1.5s before the presentation of the crucial sensory input.

One possibility is that the application of the 8-Hz lowpass filter during preprocessing may have distorted pre-component (mostly “N1” deflection) estimation. To address this possibility, we repeated the stepwise removal analysis above, but without the 8-Hz lowpass filter. For all three detection effects slightly lower *p* values (.015, .045, .265, respectively) were observed, demonstrating that our results were not driven by the lowpass filter.

## Comparison between visual and auditory fP3

Having established that the fP3 is prospectively associated with upcoming lapses of attention in both the visual and auditory domains, we asked about the extent to which visual and auditory fP3s were driven by identical intracranial dipole configurations. Because different dipole configurations should result in different scalp topographies, we conducted a detailed comparison of visual and auditory fP3 scalp topographies (Figures 4B and 4C).

To statistically test whether the fP3’s topographical distributions differed between modalities, a 28-electrode search space was defined, covering frontal, central, and central-parietal scalp regions. Specifically, a 4 by 7 grid was defined consisting of a 4-level anteriorposterior axis (frontal, frontal-central, central, and central-parietal) and a 7-level left-to-right axis (e.g., F5-F3-F1-Fz-F2-F4-F6; FC5-FC3-FC1-FCz-FC2-FC4-FC6; same for central (C) and central-parietal (CP)). Next, for each electrode and for each participant with both valid visual and valid auditory data (*n*=39), fP3s were quantified as in the analyses above (average value between 282-332ms latency for visual, 129-207ms latency for auditory). These values were averaged across 5 PT-x levels and subjected to average-reference transformation (demeaning across the 28 selected electrodes). We used an average-reference transformation to prevent sheer amplitude differences to show up as artificial interactions between electrode-location and the effect of modality (McCarthy & Woods, 1985). This allowed for a cleaner test of the interaction terms of interest and thus characterize the topographies of the visual and auditory fP3.

We then compared fP3 amplitudes with a 2 (Modality) x 2 (Detection) x 4 (Anteriority) x 7 (Left-Right) repeated-measures ANOVA. In this design, multivariate significance was only concluded when a topographical factor interacted with both the Modality effect and the Modality and Detection effects; this ensures that topographical differences between visual and auditory are paralleled by a similar topographical difference in the Detection effect. Note that in this analysis main effects of Modality, Detection and their interaction were ignored, as these effects became near-zero due to demeaning.

As can be seen in Figure 6, the visual fP3, as well as the Detection effect, was most pronounced at frontal(-central) electrodes. By contrast, for the auditory fP3, the centers of gravity were shifted more to central regions. The complete pattern was reflected in interactions between the effects of Modality, Detection, and Anteriority (3-way: *F*(3,36) = 3.5, *p* < .05), and between effects of Modality and Anteriority (2-way: *F*(3,36) = 17.0, *p* < .001). The two effects involving Modality, Detection, and Left-Right were not significant (both *F*s < .56; *p*s > .89).

Together, these results indicate that different varieties of the fP3 may exist. We here demonstrated that the visually evoked fP3 is relatively slow and frontally focused, while the auditory fP3 has an earlier onset and is focused more centrally. This may point to distinct intracranial dipole generators across modalities (see Discussion).

## Discussion

We here investigated whether the frontal P300 (fP3) tracked imminent lapses of sustained attention. We first found that visual fP3 amplitudes reveal imminent hits and misses already 1.5s before target detection was possible. In Experiment 2, we demonstrated the robustness of this effect, and crucially extended this effect from the visual to the auditory domain. Our results demonstrate that the fP3 is a supramodal marker of imminent lapses of attention.

### fP3 amplitudes as a supramodal marker of imminent attentional lapses

In the present work, the prospective association between frontal P3 (fP3) and subsequent behavioral-target detection was replicated, again confirming the validity of the use of the visual fP3 as an index of susceptibility or sensitivity to potential critical events in the near future. Specifically, this was observed in a continuous temporal expectancy task (CTET; O’Connell et al., 2009), where fP3s were elicited by non-target frames during the few seconds preceding a target frame that had to be behaviorally responded to. Leveraging the relatively large sample size in the current experiment, we demonstrated that this association was already reliable 4s before the crucial incoming visual input. That is, on average, an fP3 elicited 4 seconds preceding a detection target was correlated with whether that visual target was detected or not.

The prospective association between fP3 and subsequent target detection also holds for the auditory domain, although for a shorter time span than it does for the visual modality (again 1.5 second; note that the sample size was slightly lower, 40 versus 46, for the auditory compared to the visual task version). This is of considerable theoretical relevance, given that the original claims about fP3 as an index for susceptibility were formulated in the context of auditory probes in a context of visual tasks (Vander Heiden et al., 2018, 2020, 2022; Wester et al., 2008).

It remains to be seen to what extent these prospective relations would hold as a real prediction at the level of individuals and individual trials. One fruitful way forward could be the implementation of machine-learning based classification techniques (e.g., Chidharom et al., 2025), which could benefit from the present insights to provide a spatial-temporal template for the fP3 – although optimal results may be achieved when adjusting such templates separately for each individual.

While individual prediction may have practical implications in, e.g., process-control contexts, the present findings also bear theoretical relevance. It has been proposed (O’Connell et al., 2009) that fP3 in the CTET context can be viewed as a mechanism that tracks the temporal structure of the task. As noted, others have advocated an interpretation of fP3 in terms of a general behavioral interrupt, resulting in a state of general motor inhibition or neural freezing, in response to an unexpected event that warrants scrutiny before any action is initiated (Kenemans, 2015; Polich, 2007; Wessel & Aron, 2013, 2017). It may be interesting to integrate these theoretical notions. The idea of a temporal-tracking mechanism was especially inspired by the observation that fP3 was also elicited by the longer duration of the target frame, i.e., at some 1100ms after pre-target onset, and without any change in stimulus (O’Connell et al., 2009). This suggests that fP3 can also be elicited by an internal event, i.e., the detection of the absence of a stimulus. In turn, the fP3s elicited by preceding non-target frames could reflect the anticipation of this stimulus absence, the anticipation leading to transient neural freezing after each non-target frame. As noted by O’Connell et al. (2009, p.8611), this anticipatory response may be relatively disengaged during the seconds preceding a failed detection.

Reduced anticipatory fP3s preceding failed detections may reflect a more global transient dysfunction of multiple brain as well as peripheral processes, as demonstrated recently in the context of failed or slowed detections (also termed “attentional lapses”) in relation to sleep deprivation (Yang et al., 2025). Specifically, the involvement of pupil constriction (Aston-Jones & Cohen, 2005; Strauch et al., 2022) during failed-detection episodes suggests the involvement of drops in noradrenergic activity. In turn, this may result from an exhausting effect of sleep deprivation on noradrenergic activity, because the noradrenergic system is relatively silent during normal sleep. It is conceivable throughout that transient drops in noradrenergic activity can also occur without dedicated manipulations of sleep quality preceding task performance. Evidence for this comes from studies demonstrating that pupil size tracks fluctuations in task performance (Koevoet et al., 2023, 2024; Robison & Unsworth, 2019) and can be used to track upcoming lapses of attention (Keene et al., 2022). In this context, it is worthwhile to note that the relation between reduced fP3s and failed detections cannot be attributed to increasing anticipation of a to be detected target, because overall fP3s did not become smaller with longer time intervals preceding an eventual target frame.

### Similarities and differences between the visual and auditory fP3

Having established the prospective association between fP3 amplitudes and imminent detection of visual and auditory targets, we examined whether visually- and auditorily-evoked fP3s were intracranially generated in a similar fashion. Our detailed topographical comparison (Figure 6) suggested that the fP3 may come in different varieties: Whereas the auditory fP3 has a focus more on central scalp regions, the visual fP3 is more frontally pronounced. This could reflect a difference in the underlying configuration of intracranial sources: Equivalent dipoles may be located or oriented differently, or both, for visual versus auditory task stimuli. This in turn may be related to a difference in connections to, or from, other brain regions to the presumably dorsal-medial-frontal source region of the fP3. One possibility is that these may involve the contribution of visuo-motor neurons (Raos, Umilta, Murata, Fogassi, & Gallese, 2006), which may differ for visual compared to auditory contexts.

By the same token, the well-established novelty fP3 may have characteristics that may differ in similarly subtle ways from the presently examined CTET fP3s. In contrast to the currently employed CTET tasks, where targets and non-targets are presented within the same modality, most novelty fP3 studies use auditory novels that are essentially task-irrelevant probes in the context of a (mostly visual) task. Indeed, the novelty fP3 is in fact augmented by interspersing the novels in an auditory stream in which other auditory stimuli have behavioral relevance (Van der Heiden et al., 2018; Wester et al, 2007). Additionally, whereas novels are mostly task-irrelevant, they are also mostly unique in the sense of hardly being repeated; in contrast, the CTET presents hundreds of identical non-target frames, which are however task-relevant; the very fact of them eliciting robust fP3s is consistent with the effect of global task relevance on the novelty fP3. Future work is necessary to chart the similarities and differences between the here observed fP3s and those in novelty fP3 studies.

Another case in point concerns the peak latency of fP3 relative to the eliciting stimulus. For both task-relevant repeated visual non-target frames and for task-irrelevant unique auditory novels, fP3 latency amounts to about 300ms; but for task-relevant repeated auditory nontarget frames the latency is much shorter, at about 170ms. This is reminiscent of modality effects on the latency of the fP3 as commonly observed in stop-signal paradigms: Whereas the stop-related fP3 for auditory stop signals peaks well before 200ms, for visual stop signals it does so after 200ms (Kenemans et al., 2023).

One may argue that without a proper magnitude estimation procedure such inter-modality differences are hard to interpret. What remains conspicuous however, is the overall shorter latencies of stop-signal fP3 relative to auditory novels and visual CTET frames. As mentioned already briefly, a detailed analysis based on a combination of novelty and stop-signal paradigms suggested that the fP3s elicited in both contexts can be viewed as a unitary process (Wessel & Aron, 2013). Note that in that study, the fP3 was defined in terms of independent components that were based on scalp topography, frequency content, and correlation with stopping success. In turn, such a unitary interpretation does fit notions about fP3 as a reflecting a state of general motor inhibition, which could be especially instrumental in response to a stop signal.

In all, additional detailed spatial-temporal comparisons between potentially different fP3 varieties seem to be worthwhile. They could be realized using independent-component analysis (Makeig et al., 1995), or by using other approaches in which topographical, temporal, and frequency characteristics are combined. In turn, the results of such analyses could be further informed by detailed explorations of the functional neuroanatomy of dorsal-medial frontal cortex (e.g., the presupplementary motor area).

To conclude, our results provide evidence that the EEG-frontal P3 wave reflects susceptibility to incoming sensory input. Specifically, we demonstrated that fP3 amplitudes tracked upcoming lapses of attention. We extended previous work by showed that this effect holds not only for the visual but also for auditory domain. Our findings were found in a context in which the timing of these events is quite unpredictable. Although the spatial-temporal characteristics of the visual and auditory fP3s are robustly different they both have a dorsalmedial frontal scalp signature, and they both reliably track imminent lapses of sustained attention.

## Supplementary materials

**Table 1.** Full repeated-measures ANOVA output for Experiment 1 (fP3 Visual).

| **Effect** | ***F*** | ***P*** | **$\eta_p^2$** |
| --- | --- | --- | --- |
| Detection | 10.75 | .003 | .31 |
| Latency | 52.85 | <.001 | .69 |
| Pre-target interval | 3.62 | .02 | .41 |
| Detection * Latency | 0.52 | .48 | .02 |
| Detection * Pre-target interval | 1.01 | .42 | .16 |
| Latency * Pre-target interval | 1.15 | .36 | .18 |
| Detection * Pre-target interval * Latency | 0.92 | .47 | .15 |

**Table 2.** Full repeated-measures ANOVA output for Experiment 2 (fP3 Visual).

| **Effect** | ***F*** | ***P*** | **$\eta_p^2$** |
| --- | --- | --- | --- |
| Detection | 26.45 | < .001 | .37 |
| Latency | 148.18 | <.001 | .77 |
| Pre-target interval | 1.53 | .21 | .13 |
| Detection * Latency | 14.47 | <.001 | .24 |
| Detection * Pre-target interval | 0.42 | .79 | .04 |
| Latency * Pre-target interval | 1.44 | .24 | .12 |
| Detection * Pre-target interval * Latency | 1.15 | .35 | .10 |

**Table 3.** Full repeated-measures ANOVA output for Experiment 2 (fP3 Auditory).

| **Effect** | ***F*** | ***P*** | **$\eta_p^2$** |
| --- | --- | --- | --- |
| Detection | 6.17 | .02 | .14 |
| Pre-target interval | 0.42 | .79 | .05 |
| Detection * Pre-target interval | 1.03 | .41 | .10 |

